# Chemical map–based prediction of nucleosome positioning using the Bioconductor package nuCpos

**DOI:** 10.1101/2019.12.25.888305

**Authors:** Hiroaki Kato, Mitsuhiro Shimizu, Takeshi Urano

## Abstract

**Background:** Assessing the nucleosome-forming potential of specific DNA sequences is important for understanding complex chromatin organization. Methods for predicting nucleosome positioning include bioinformatics and biophysical approaches. An advantage of bioinformatics methods, which are based on *in vivo* nucleosome maps, is the use of natural sequences that may contain previously unknown elements involved in nucleosome positioning *in vivo*. The accuracy of such prediction attempts reflects the genomic coordinate resolution of the nucleosome maps applied. Nucleosome maps are constructed using micrococcal nuclease digestion followed by high-throughput sequencing (MNase-seq). However, as MNase has a strong preference for A/T-rich sequences, MNase-seq may not be appropriate for this purpose. In addition to MNase-seq–based maps, base pair–resolution chemical maps of *in vivo* nucleosomes from three different species (budding and fission yeasts, and mice) are currently available. However, these chemical maps have yet to be integrated into publicly available computational methods.

**Results:** We developed a Bioconductor package (named nuCpos) to demonstrate the superiority of chemical maps in predicting nucleosome positioning. The accuracy of chemical map–based prediction in rotational settings was higher than that of the previously developed MNase-seq–based approach. With our method, predicted nucleosome occupancy reasonably matched *in vivo* observations and was not affected by A/T nucleotide frequency. Effects of genetic alterations on nucleosome positioning that had been observed in living yeast cells could also be predicted. nuCpos calculates individual histone binding affinity (HBA) scores for given 147-bp sequences to examine their suitability for nucleosome formation. We also established local HBA as a new parameter to predict nucleosome formation, which was calculated for 13 overlapping nucleosomal DNA subsequences. HBA and local HBA scores for various sequences agreed well with previous *in vitro* and *in vivo* studies. Furthermore, our results suggest that nucleosomal subsegments that are disfavored in different rotational settings contribute to the defined positioning of nucleosomes.

**Conclusions:** Our results demonstrate that chemical map–based statistical models are beneficial for studying nucleosomal DNA features. Studies employing nuCpos software can enhance understanding of chromatin regulation and the interpretation of genetic alterations and facilitate the design of artificial sequences.

## Background

The nucleosome is a structure in which a ~147-base pair (bp) double-stranded DNA forms a left-handed super-helix to wrap a histone octamer consisting of two copies of histones H2A, H2B, H3, and H4 [1–3]. The positioning of nucleosomes is defined by rotational and translational settings [4, 5]. Rotational settings define the histone-DNA contact surfaces on the DNA string, whereas translational position is the distinct position of the nucleosome (i.e., position of the dyad base) with respect to the genomic coordinate. The positioning of nucleosomes along genomic DNA is critical for various biological events, including transcription, DNA replication and repair, and chromosome segregation [4]. In these events, nucleosome positioning is combinatorially regulated by *trans*-acting factors (e.g., transcription factors and chromatin remodeling factors) and by the inherent suitability of the DNA sequence for nucleosome formation [5–7]. Focusing on the latter requirement, computational methods that assess the nucleosome-forming potential of given DNA sequences have been developed [8].

These methods are classified into two groups: biophysical and bioinformatics [8, 9]. Most biophysical methods are based on the structures of reconstituted nucleosome core particles (NCPs) such as NCP147, which contains a palindromic 147-bp sequence with intrinsic positional stability [3, 10, 11]. The use of the palindromic sequence optimized for homogeneous nucleosome reconstitution and the highest resolution of the structure contributed greatly to the development of prediction algorithms. However, relying on the representative structure itself could also be a potential weakness, as there could be unrecognized natural intra-nucleosomal DNA elements that mediate specific physiologic functions in cells.

In contrast to the biophysical prediction methods, bioinformatics methods are based on maps of *in vivo* nucleosomes determined using high-throughput sequencing techniques. The use of complex, natural nucleosomal sequences may cause difficulty in the interpretation of prediction results; however, it may also lead to unexpected findings that enhance understanding of nucleosome-based regulation.

A weakness of currently available bioinformatics methods is that the statistical models are constructed based on *in vivo* nucleosome maps generated using micrococcal nuclease (MNase) digestion followed by high-throughput sequencing (MNase-seq). MNase digests linker DNA that connects NCPs, allowing researchers to obtain nucleosomal DNA that has been protected from enzymatic digestion. In MNase-seq experiments, DNA fragments of mono-nucleosome size (~150 bp) are sequenced and mapped to the reference genome. However, as MNase strongly prefers A/T-rich sequences, unwanted digestion of A/T-rich nucleosomal sequences and less-effective digestion of G/C-rich linkers may occur, leading to variation in nucleosome fragment size [7, 12]. Because of this enzymatic bias, precise determination of the genomic coordinates of *in vivo* nucleosomes using MNase-seq (i.e., dyad base calling) is challenging. In addition, although some correction methods for MNase-seq coverage against enzymatic bias have been developed [13–15], no method that corrects the deduced nucleosome position by shifting it to an appropriate rotational position is available. Thus, the resolution of MNase-seq–based maps with respect to the genomic coordinates is expected to be inevitably low.

One bioinformatics method considered successful in a thorough comparative study [9] is NuPoP, which is built upon a duration hidden Markov model (dHMM) that considers information regarding both the nucleosome and linker DNA [16]. NuPoP enables the prediction of nucleosome positioning in nine different species by rescaling the base composition of both the nucleosome and linker models for the budding yeast *Saccharomyces cerevisiae* with those of target organisms. The budding yeast models are generated using an MNase-seq–based nucleosome map by considering the central base of each nucleosomal fragment of varying length the dyad base [16]. This may particularly affect construction of the time-dependent statistical model for the nucleosome state, which ideally requires that the dyad bases be determined at bp resolution. The construction of the linker model, which requires extraction of linker sequences as genomic sequences not covered by nucleosomes, may also be affected. Thus, although selecting the MNase-seq–based map was the best practice at the time of the software’s development, the prediction results output provided by NuPoP can be biased by both the enzymatic preference of MNase and the difficulty of dyad base calling.

A breakthrough in nucleosome mapping occurred with the development of a method for site-directed chemical cleavage of nucleosomal DNA followed by high-throughput sequencing [17, 18]. This technique, called chemical mapping, was first applied to budding yeast and subsequently to the fission yeast *Schizosaccharomyces pombe* and embryonic stem cells of the house mouse, *Mus musculus* [17, 19–22]. In cells expressing histone H4 with an S47C amino acid substitution, nucleosomal DNA is specifically cleaved near the nucleosome center by site-directed chemical cleavage. By examining the cleavage sites, previous studies determined the dyad positions of *in vivo* nucleosomes in the three species at bp resolution [17, 19, 22]. The published maps contained whole and representative sets of identified nucleosomes, which are referred to as “redundant” and “unique” nucleosomes, respectively [17, 19, 22]. In addition to the histone H4-S47C approach, chemical mapping of nucleosomes using a histone H3 mutation (Q85C) determined the precise positions of −1 and +1 nucleosomes for protein-coding genes [12]. However, the bp-resolution chemical maps have not yet been implemented in publicly available software for prediction of nucleosome positioning.

One goal of nucleosome positioning prediction is *in silico* reproduction of *in vivo* observations, preferably at the locus level. If this goal is achieved, predicting the effects of genetic alterations on nucleosome positioning and prediction-based engineering of nucleosome-forming and -depleted sequences would also be practical. In this regard, budding and fission yeasts would serve as preferable model organisms, because nucleosomes formed on native and modified sequences *in vivo* have been precisely mapped [23–28]. These nucleosomes are suitable for *in silico* reproduction of *in vivo* observations. Importantly, for the currently available bioinformatics method NuPoP, the reliability of locus-level nucleosome prediction has seldom been demonstrated, despite its wide use in previous studies [29–38].

In this study, we developed a nucleosome positioning prediction method that utilizes available chemical maps of three species for construction of dHMMs. We show that the prediction accuracy of chemical map–based models is higher with respect to the genomic coordinates than that of MNase-seq–based models. The chemical map–based models appear to recognize appropriate sides of DNA strings as histone-DNA contact surfaces. The effects of genetic alterations on nucleosome positioning previously observed in living yeast cells were reasonably reproduced *in silico*, suggesting that this software is useful for chromatin engineering and interpretations of mutation effects. We found that the prediction accuracy for mice was lower than that for yeasts, which should be acknowledged by software users. Our software, nuCpos, is available at the Bioconductor website (https://doi.org/doi:10.18129/B9.bioc.nuCpos).

## Results

### Chemical map–based prediction of nucleosome positioning in yeasts

In order to determine whether the bp-resolution chemical maps are suitable for predicting nucleosome positioning, we trained dHMMs with the nucleosome maps generated with H4-S47C–dependent DNA cleavage (see **Methods** and **Table 1**). The core dHMM algorithm implemented in our method (nuCpos) was not altered from the original algorithm developed for the MNase-seq–based method (NuPoP) to clarify the advantages of using the chemical maps.

**Table 1.**
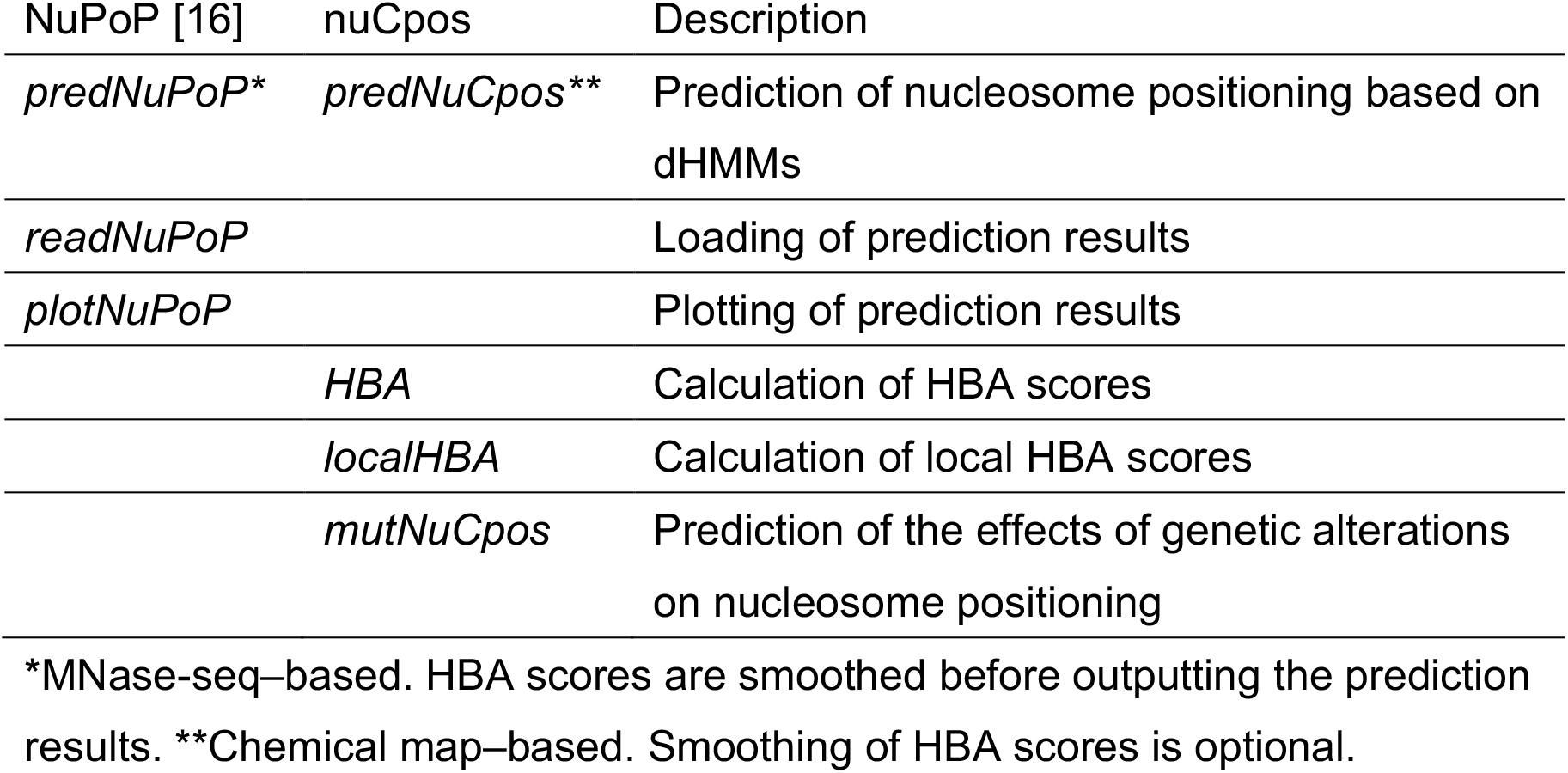
Functions of the software packages used in this study.

First, we predicted nucleosome positioning in the genomes of budding and fission yeasts (denoted ‘Sc’ and ‘Sp’, respectively, in **Figure 1**) with models that were constructed with *in vivo* nucleosome maps of target species. Aiming to evaluate the prediction accuracy with respect to the genomic coordinates, prediction results were assessed by checking the matching status of predicted nucleosomes with those experimentally determined by means of histone H4-S47C–based chemical cleavage [17, 19]. Here, MNase-seq–based maps are inappropriate to use as true-position references due to their lower resolution, as described in the Background section. Ideally, prediction results should be evaluated at 1-bp resolution, so that it will be possible to accurately discern the rotational settings of given nucleosomes. As the resolution of the chemical maps that we used for model construction could be 1-2 bp or lower, the resolution of the prediction results would be expected to be at a similar level. Indeed, the distance between chemically predicted nucleosomes and the nearest *in vivo* nucleosomes was generally 0 bp (located at the same position) or 1 bp (located at the juxtaposed nucleotide positions) (**Figure 1A**, right panel).

**Figure 1.**
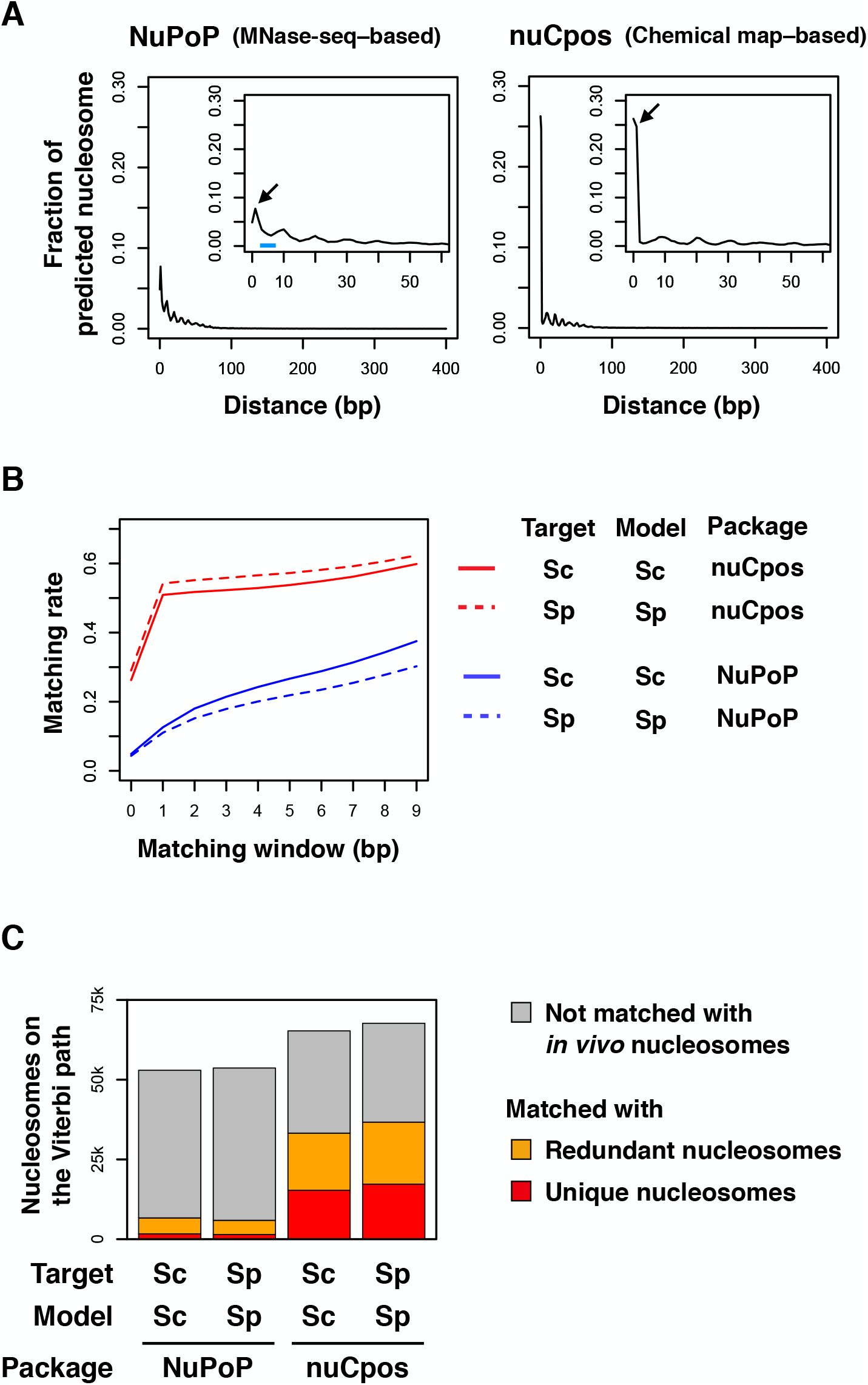
Comparison of prediction accuracy between chemical map–based and MNase-seq–based models for yeasts. (**A**) Distance between predicted nucleosomes and nearest *in vivo* nucleosomes. Fraction of predicted nucleosomes on the Viterbi path located at a particular distance from the nearest *in vivo* nucleosomes is plotted. Species of the target genome and for model construction were both *S. cerevisiae*. Magnified view is presented inside each plot. Arrows point to the fractions of predicted nucleosomes that are 1-bp apart from *in vivo* nucleosomes. (**B**) Matching rate between predicted and in vivo nucleosomes with variable matching windows. The species of genomic sequences used for prediction (Target) and the species of the chemical map used for model construction (Model) are indicated. “Sc” stands for *S. cerevisiae*, whereas “Sp” stands for *S. pombe*. Indicated R packages were used for predictions. (**C**) Matching of nucleosomes on the Viterbi paths with *in vivo* unique and redundant nucleosomes at 2-bp resolution. Note that the redundant nucleosome datasets contain both unique nucleosomes (red), which are representative, and non-representative nucleosomes (orange).

In order to determine the appropriate resolution for examining prediction results, we calculated the matching rate between predicted and *in vivo* nucleosomes with stepwise widening of the matching window (**Figure 1B**). Here, we defined the matching window as the allowable distance between nucleosomes regarded as matched. In chemical map–based predictions (nuCpos), the matching rate doubled to over 50% when the matching window was widened from 0 bp (1-bp resolution) to 1 bp (2-bp resolution) (**Figure 1B**, red lines). However, the rate did not markedly increase upon further widening of the matching window, suggesting that the chemical models distinguish the histone-facing side of the DNA string.

In MNase-seq–based predictions (NuPoP), by contrast, the matching rate increased gradually as the matching window was widened from 0 to 9 bp (**Figure 1B**, blue lines). The matching rate drew a very loose anti-S shape, suggesting that the MNase-seq–based model still recognized the histone-facing side of the DNA string, but only to a small extent. We also noticed that a substantial population of nucleosomes predicted with the MNase-seq–based model was positioned approximately 5 nucleotides away from the *in vivo* nucleosomes (**Figure 1A**, left panel, marked with a blue horizontal bar). These observations suggest that the MNase-seq–based model has difficulty in discriminating rotationally mispredicted nucleosomes. Considering these results, we concluded that prediction results should be evaluated at 2-bp resolution, as further widening of the matching window would increase the misprediction rate.

Figure 1C presents the evaluation of prediction results against representative “unique” and whole “redundant” nucleosomes [17, 19] at 2-bp resolution. When the budding yeast MNase-seq–based model (NuPoP) was applied to the prediction of nucleosome positioning in the budding yeast genome (Target: Sc, Model: Sc), only 3.2% and 12.6% of the nucleosomes on the most probable (Viterbi) path matched *in vivo* unique (red) and redundant (orange) nucleosomes, respectively. When the fission yeast model was applied to the prediction of nucleosome positioning in the fission yeast genome (Target: Sp, Model: Sp), the MNase-seq–based model scored 2.7% and 11.0%, respectively. In contrast, when the budding yeast chemical map–based model (nuCpos) was used for predictions (Target: Sc, Model: Sc), 23.5% and 50.9% of the nucleosomes on the Viterbi path matched unique and redundant nucleosomes, respectively. For the fission yeast (Target: Sp, Model: Sp), the corresponding scores were 25.5% and 54.2%, respectively. Thus, with the chemical map– based models, over half of the nucleosomes predicted as being on the Viterbi path are present in living cells at exactly the same positions or at the juxtaposed nucleotide positions.

In the chemical maps of budding and fission yeasts, *in vivo* unique nucleosomes accounted for 19.6% and 17.8%, respectively, of all redundant nucleosomes [17, 19]. Thus, in the MNase-seq–based predictions, enrichment of unique nucleosomes on the Viterbi paths was only 1.3- and 1.4-fold, respectively. In contrast, the chemical map–based predictions exhibited 2.4- and 2.6-fold enrichment, respectively. The higher enrichments suggest that chemical map–based models better distinguish sequences favorable for nucleosome formation [17, 19], compared with MNase-seq–based models.

Since the dHMMs were constructed using nucleosome and linker information based solely on the unique nucleosomes (see **Methods**), detection of portion of the redundant nucleosomes (from which unique nucleosomes have been removed) is worthwhile. The nucleosomes in this category accounted for 27.4% and 28.7% of the Viterbi nucleosomes in the chemical-map–based predictions for the budding and fission yeasts, respectively, and the proportions were higher than the MNase-seq–based predictions (9.4% and 8.3%, respectively). Giving these results, the chemical map–based models provide better prediction outcomes with respect to the genomic coordinates than the MNase-seq–based models.

### Prediction of *in vitro* reconstituted nucleosome positions

The evaluations described above suggested that chemical map–based models are superior in terms of the prediction of rotational settings. To confirm this observation, we compared HBA scores [16] calculated along the original 282-bp Widom 601 sequence [10] with the chemical map–based and MNase-seq– based models (**Figure 2**). HBA scores are calculated in the dHMM algorithms in order to examine the suitability of given 147-bp sequences for nucleosome formation [16], and they are expected to predict the rotational setting of nucleosomes if the resolution of the nucleosome map used for model construction is sufficiently high. The original 282-bp Widom 601 sequence contained primer sequences for the SELEX study at both ends and additional sequences flanking the nucleosome-forming 147-bp sequence centered at nucleotide position 154 [10]. When analyzed using the chemical map–based model, discrete HBA peaks with an interval of approximately 10 bp were observed at positions 124, 135, 143, 154, 164, and 175 (asterisks). This periodicity indicated that nearly the same surface of the DNA string was predicted to be in contact with the histone core in these potential translational positions. Among these positions, the one with the highest HBA score corresponds to the translational position of the *in vitro* stable nucleosome (position 154, orange vertical lines). In contrast, the MNase-seq–based HBA score was relatively low at position 154, and the peaks did not exhibit the apparent 10-bp periodicity. Analyses of the mouse mammary tumor virus (MMTV) 3’–long terminal repeat (LTR) sequence and *Xenopus borealis* 5S RNA gene also showed the advantage of the chemical model (**Additional file 1: Figure S1**). Collectively, the chemical map–based model reliably predicted the rotational settings of the nucleosomes reconstituted with these suitable sequences.

**Figure 2.**
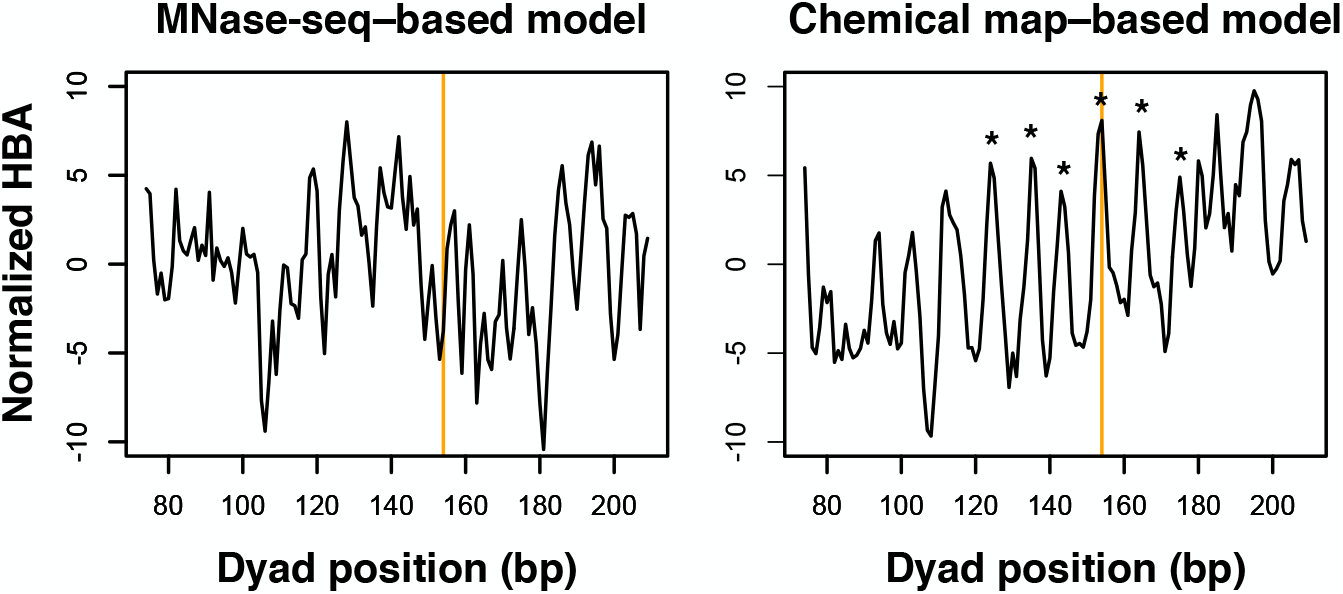
HBA scores along the Widom 601 sequence. HBA scores along the 282-bp original Widom sequence were calculated using chemical map–based and MNase-seq–based budding yeast models. The scores were normalized by subtracting the mean value from the raw value on a per sequence basis. The HBA score for a given 147-bp nucleosome sequence was assigned to its dyad position. Orange vertical lines indicate the dyad positions of *in vitro* reconstituted nucleosome: nucleotide position 154. Asterisks indicate high-scoring HBA positions around the *in vitro* positioning site.

### Prediction of *in vivo* nucleosome positions

Next, we calculated HBA scores around −1 and +1 nucleosomes for 5,542 protein-coding genes identified by means of histone H3-Q85C–based chemical cleavage [12]. Due to the differences in detection methods, positions of H3-Q85C–based nucleosomes are not always identical to those of the corresponding H4-S47C–based nucleosomes [12]. In this regard, the H3-Q85C–based map differed from the H4-S47C–based map that was used for the statistical model construction. Thus, the H3-Q85C–based nucleosomes would be a good reference for evaluation of prediction accuracy at bp-resolution.

As shown in **Figure 3A**, mean HBA scores calculated using both models exhibited a periodicity with a 10-bp interval. However, the amplitude of the chemical map–based HBA scores (red) around the tested nucleosomes was larger than that of the MNase-seq–based HBA scores (blue). This suggested that the chemical model could clearly distinguish the surface of the DNA string that interacts with the histone core even when natural genomic sequences were queried. We found that the chemical map–based HBA scores were highest at the −1 and +1 nucleosome positions (0 bp). Chemical map–based HBA scores for the three neighboring translational positions (10, 20, and 30 bp in distance) relative to the nucleosome-depleted regions (marked with asterisks in **Figure 3A**) remained relatively high despite the gradual increase in the A/T frequency of the tested sequences (**Figure 3B**). After the A/T frequency reached a plateau (A/T≈0.64), the chemical map–based HBA scores began to decrease. In contrast, MNase-seq–based HBA scores at the −1 and +1 nucleosome positions (0 bp) were already at moderate levels, and they simply decreased as the A/T frequency increased. Thus, the MNase-seq–based model was adversely influenced by the enzymatic bias of MNase, whereas the chemical map–based model was not.

**Figure 3.**
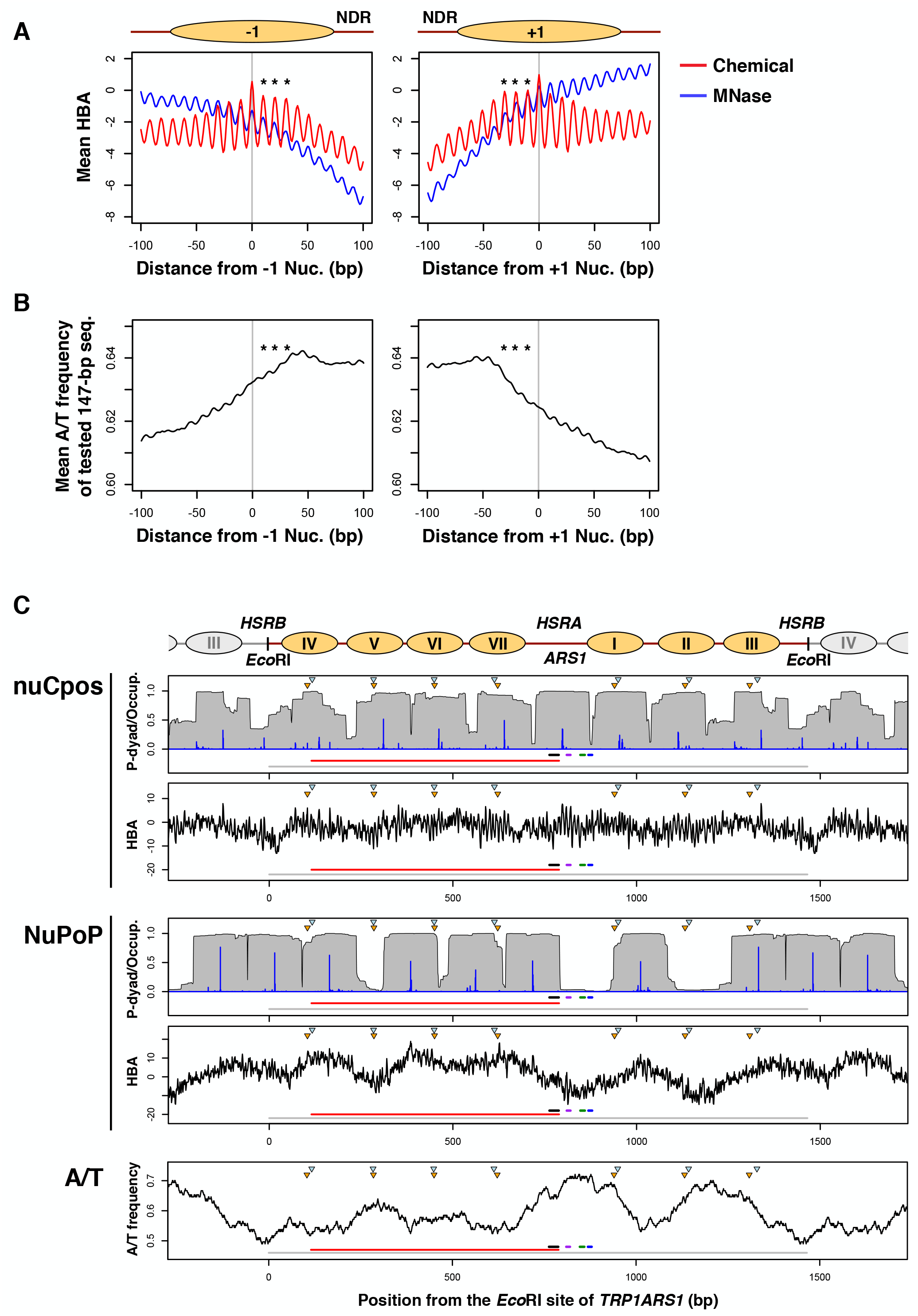
Comparison between chemical map–based and MNase-seq– based models on selected *in vivo* regions. (**A and B**) Chemical map–based and MNase-seq–based HBA scores (**A**) and A/T nucleotide frequency (**B**) along the sequences containing budding yeast −1 and +1 nucleosomes for protein-coding genes. Gray vertical lines indicate the dyad positions of respective nucleosomes. Nucleosomes of which the dyad base is located at 0 bp are shown as ovals above the plots. Asterisks indicate positions with relatively high HBA scores. NDR: nucleosome-depleted region. (**C**) Prediction results for the budding yeast *TRP1ARS1* mini-chromosome. Schematic representation of *in vivo* nucleosome positioning is shown above the plots. Nucleosomes labeled I through VII are indicated as ovals. Note that this sequence is circularized *in vivo* by being linked at the *Eco*RI sites. The top two panels show the prediction results output by nuCpos, whereas the next two panels show NuPoP results. The upper panel in each set shows predicted occupancy of the nucleosome (Occup., gray polygons) and probabilities of the tested 147-bp sequences for being in the nucleosome state (P-dyad, blue vertical lines). The lower panels show HBA values for the tested 147-bp sequences calculated using the indicated models. The very bottom panel shows the A/T-frequencies for the tested 147-bp sequences. Horizontal lines at the bottom of each plot indicate a unit of the circular *TRP1ARS1* mini-chromosome (colored in gray, nucleotide positions 0 to 1,464), the coding region of *TRP1* (red, 115-879), and the *B3* (black, 763-788), *B2* (purple, 810-820), *B1* (green, 847-859), and *ACS* (blue, 869-879) elements of *ARS1*. Inverted triangles indicate the histone H4 S47C-dependent cleavage centers determined by indirect end-labeling; orange for mini-chromosome and light blue for genomic experiments [27].

The higher accuracy of chemical map–based models at the genomic level prompted us to perform a locus-level evaluation of prediction results, using the budding yeast *TRP1ARS1* minichromosome [25, 27]. On this well-studied, circular, 1,465-bp DNA, three (numbered I to III) and four (IV to VII) nucleosomes are interspersed with two nucleosome-depleted regions or nuclease-hypersensitive regions designated *HSRA* and *HSRB*. *HSRA* includes the DNA replication origin *ARS1*, whereas *HSRB* is located upstream of the *TRP1* marker gene. At a glance, the chemical map– and MNase-seq–based model prediction results appeared very different (**Figure 3C**). Each model suggested eight and seven representative nucleosomes, respectively. Seven of the eight nucleosomes predicted by the chemical map–based model (**Figure 3C**, nuCpos) reasonably matched the positions of *in vivo* nucleosomes determined previously [27]. The only exception was a predicted nucleosome located in *HSRA*, in which functional *ARS1* would be occupied by the ARS-binding factor Abf1 and the origin recognition complex instead of the nucleosome to maintain this circular DNA across generations [39]. This prediction was reasonable, as DNA replication origins are known to be covered by nucleosomes when the subunits of the origin recognition complex or Abf1 are perturbed *in vivo* [39–41].

In this native sequence, the chemical map–based HBA scores exhibited a periodicity of an approximately 10-bp interval (**Additional file 1: Figure S2**, nuCpos). As the dHMM considers the linker length distribution as a major factor affecting nucleosome-linker state transition and does not allow nucleosomes with shorter DNAs (<147 bp), nucleosome positions tend to be selected in a limited manner. Although there were many high-HBA positions, only a few were clearly selected by the dHMM as high-probability nucleosome dyad positions. Thus, the predicted occupancy should be cautiously considered, as it appears to provide only a rough image of chromatin state. Instead, it should be kept in mind that periodic HBA scores may contribute to redundant nucleosome positioning *in vivo*.

In contrast to the chemical map–based model, most of the nucleosome locations predicted by the MNase-seq–based model were not in agreement with their positions *in vivo* (**Figure 3C**, NuPoP); most of the *in vivo* nucleosome centers (inverted triangles in figure) were located in the predicted linkers or nucleosome ends. This result was consistent with the lower genome-wide prediction accuracy (**Figure 1**). Importantly, MNase-seq–based HBA scores for this sequence appeared to mirror the A/T-content over the X-axis (**Figure 3C**, A/T); high-HBA positions were generally A/T-poor, and low-HBA positions were A/T-rich. In addition, the local amplitude of MNase-seq–based HBA scores was apparently lower compared with the chemical map–based model (**Additional file 1: Figure S2**). These observations also demonstrated that MNase-seq– based model prediction results are strongly influenced by the A/T preference of MNase. A/T-content exhibited a similar effect in the MNase-seq–based nucleosome positioning prediction around the fission yeast *ura4*^+^ gene (**Additional file 1: Figure S3**) [42]. Given these results, we concluded that the chemical map–based models output better genome-wide and locus-level prediction results.

### Prediction of effects of genetic alterations on nucleosome positioning

Next, we examined whether the effects of genetic alterations on nucleosome positioning could be predicted. In **α** cells of budding yeast, the **a**-cell-specific gene *BAR1* is repressed by *α2*-repressor–dependent positioning of the nucleosome at its promoter, which masks the gene’s *TATA* box [23, 26]. The chemical map–based model predicted the positioning of this repressive nucleosome with high accuracy (**Figure 4**, WT). The HBA score for this translational position (0 bp), which is 82 bp away from the *α2*-operator, was very high (HBA=8.47), suggesting that repression of the *BAR1* gene is assisted by the intrinsic suitability of the promoter DNA for nucleosome formation. When a 36-bp sequence consisting of 12 repeats of CTG is inserted at the center of this nucleosome, repression of *BAR1* and nucleosome positioning are not affected *in vivo* [26]. Prediction results agreed that this insertion should not cause nucleosome depletion (**Figure 4**, CTG_12_). Other repeat sequences causing no transcriptional derepression *in vivo* [26] were also predicted to not cause nucleosome depletion (**Figure 4**, Sac_5_ and Sac_6_). It is also known that insertion of a 30-bp A-stretch or a 10-bp CG (i.e., CpG) repeat sequence inhibits *in vivo* nucleosome formation and causes derepression of *BAR1* [26]. Remarkably, the effect of these genetic alterations on nucleosome positioning was reproduced *in silico* with the chemical map–based model (**Figure 4**, A_30_ and CG_5_). We noted that the disruptive effect of fragment insertion on nucleosome positioning was more pronounced in predictions than *in vivo* observations. Insertion of a 20-bp A-stretch, which does not cause complete nucleosome depletion *in vivo*, was clearly predicted to inhibit nucleosome formation (**Figure 4**, A_20_). However, in the case of CG repeat insertion, shortening of the repeat to 8 bp, which satisfies *in vivo* nucleosome formation [26], was predicted to preserve the capability to form nucleosomes (**Figure 4**, CG_4_). We also confirmed that the inhibition of nucleosome formation by telomeric DNA insertion demonstrated in a previous study [43] was also largely predictable (**Additional file 1: Figure S4**). Thus, overall, the effects of genetic alterations on nucleosome positioning can be predicted with the chemical map–based model.

**Figure 4.**
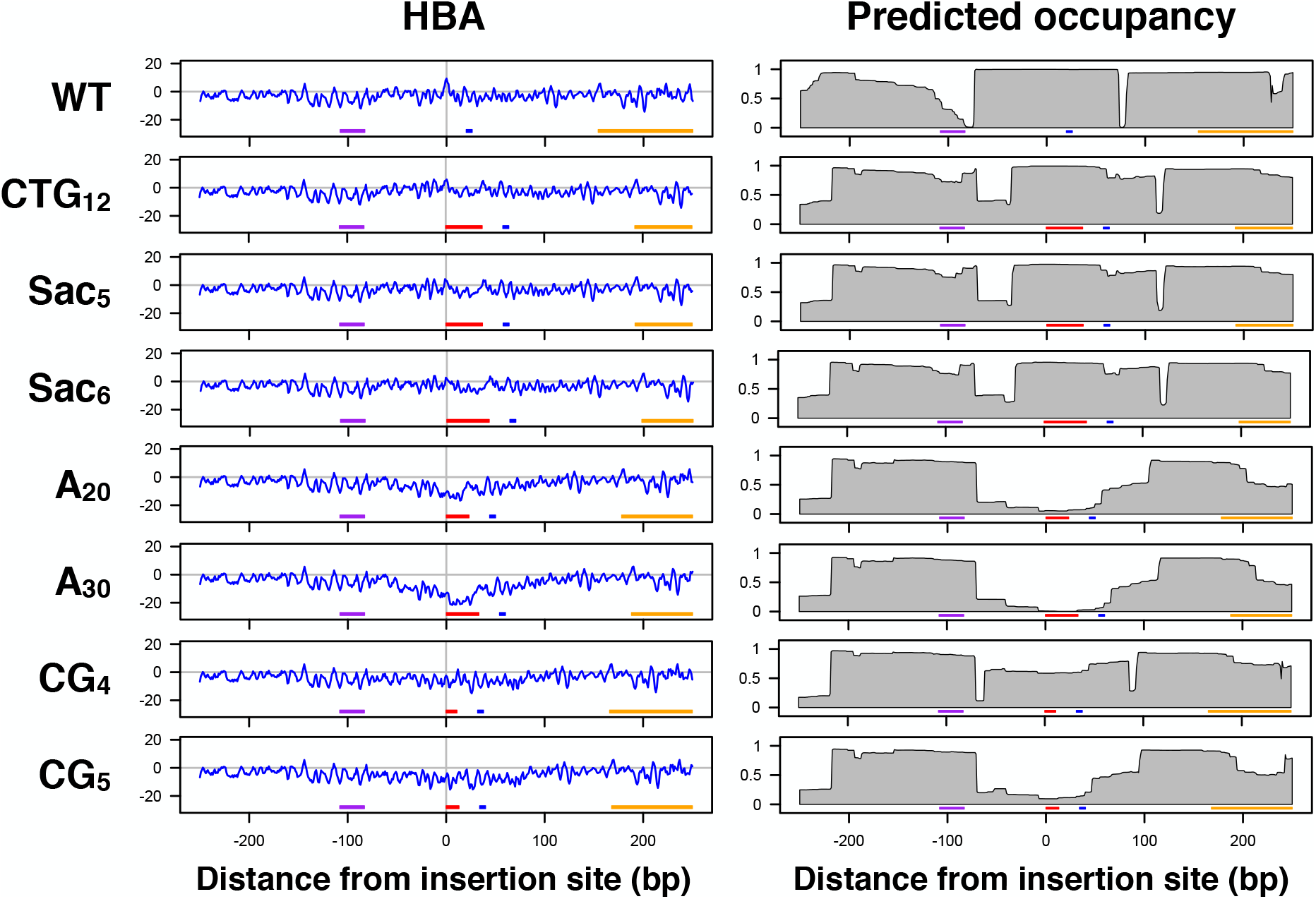
Predicting the effects of repeat insertion on nucleosome positioning. Chemical map–based HBA scores and predicted nucleosomal occupancy were calculated for original and modified *BAR1* promoter sequences. The sequences were centered at the dyad position of the repressive nucleosome, in which the indicated repeat sequences were inserted. Horizontal lines at the bottom of each plot indicate the *α2*-operator (purple), the *TATA* (blue) element, the coding region of *BAR1* (orange), and the insert (red). Note: the sequence insertion caused a shift of the left of the three predicted nucleosomes to the right to cover the *α2*-operator in the occupancy plots. Overconfidence should not be placed in this shift, as it may simply reflect that the dHMM tends to follow the linker length distribution, which generally does not allow longer linkers.

### Local HBA for nucleosomal DNA subsegments

The successful increase in prediction accuracy and better discrimination of rotationally favorable nucleosome positions with chemical map–based models suggested that the chemical map–based HBA score for a given 147-bp sequence is a good indicator of the suitability of that sequence for nucleosome formation. Considering that nucleosomal DNA makes contact with histone proteins at each superhelical location (SHL ±0.5-6.5), we defined a novel parameter, designated “local HBA”, to examine the suitability of intra-nucleosomal DNA segments for each histone-DNA contact. In order to implement this idea, a 147-bp whole nucleosomal segment was divided into 13 overlapping nucleosomal DNA subsegments, designated A through M (**Additional file 1: Figure S5**, see **Methods**). Each 20- or 21-bp segment corresponded to two histone-DNA contact sites, each of which was shared by neighboring segments. For example, segment B corresponded to SHL −5.5 and −4.5, which were shared by segments A and C, respectively. We took this overlapping approach because it remains unclear how surrounding sequences affect histone-DNA contacts. Calculating local HBA scores is conceptually the same as that of HBA calculations, which consider the probability of a whole 147-bp sequence to be a nucleosome [16]. In local HBA calculation, for each intra-nucleosomal segment of a given 147-bp DNA that wraps a histone octamer to form a nucleosome, the probability of the selected 20- or 21-bp sequence to be located at that part of the nucleosome and the probability of the same sequence functioning as a linker are calculated. The log likelihood ratio of these probabilities is defined as the local HBA score for that segment. Thus, the sum of local HBA scores for the non-overlapping set of seven 21-bp segments, A, C, E, G, I, K, and M (147 bp in total), would be nearly equal to the HBA score calculated for the same 147-bp sequence.

In order to examine whether local HBA scores indicate the suitability of nucleosomal subsegments for histone-DNA contacts, we used the 282-bp Widom 601 sequence, which was used for the above HBA analysis (**Figure 2**), as a test sequence. **Figure 5A** presents a heatmap of the calculated local HBA scores. The topmost track, marked “A”, represents local HBA scores for the A segments of 147-bp sequences centered at the indicated dyad positions. At potential translational positions with high HBA scores (**Figure 5A**, arrows), local HBA scores for some segments (e.g., segments D and E of the 147-bp sequence centered at position 154) were very high, as expected. However, each 147-bp sequence also contained segments with relatively low local HBA scores. Interestingly, at translational positions with low HBA scores (those intervened by the high-scoring HBA positions), some segments (e.g., segments F and G at position 157) exhibited very low local HBA scores. This tendency was particularly notable for the central segments E through I. The finding of these low-scoring local HBA segments suggests that sub-sequences that are disfavored in other rotational settings play a role in determining rotational settings.

**Figure 5.**
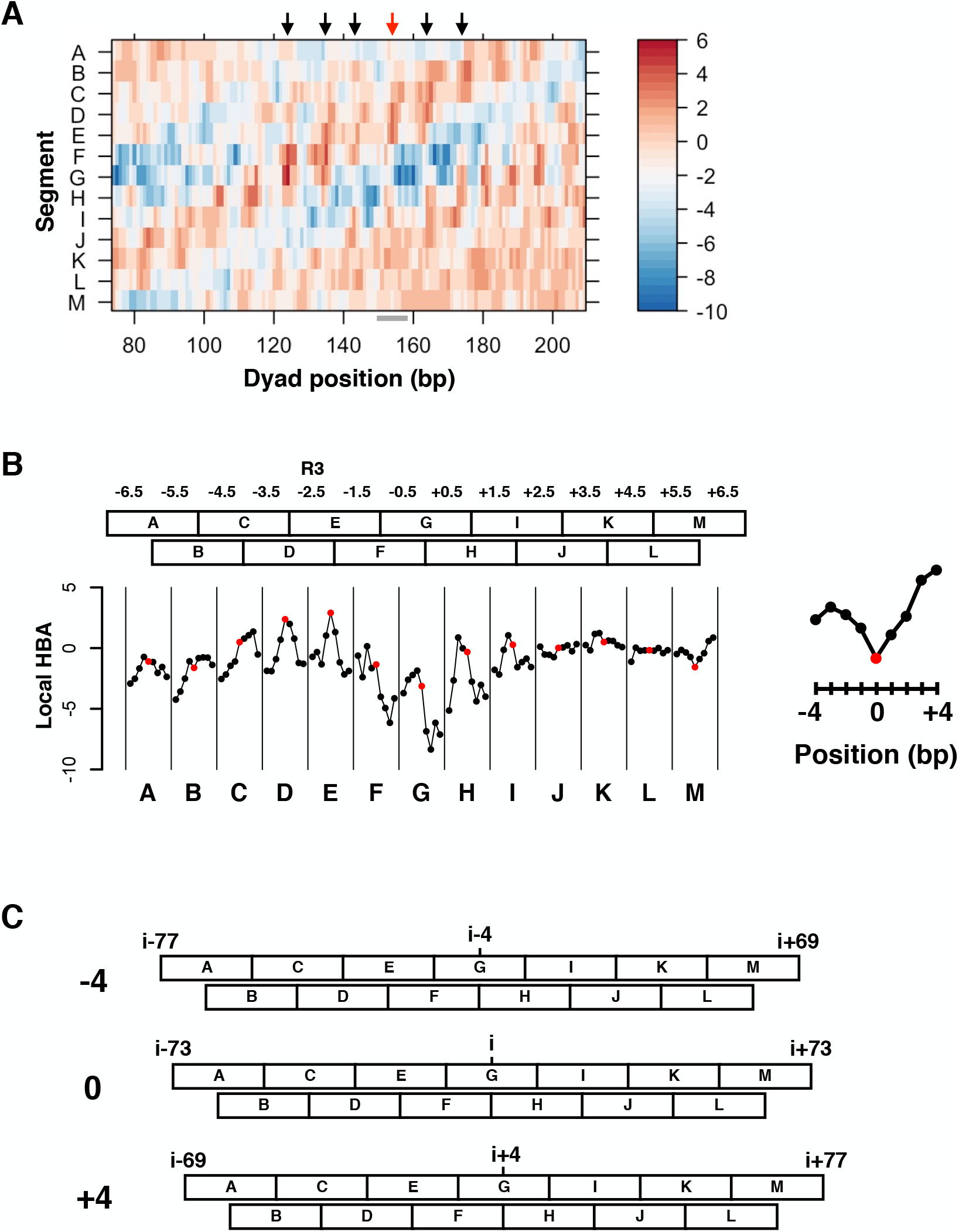
Local HBA analysis of the Widom 601 sequence. (**A**) Chemical map–based local HBA scores along the 282-bp Widom 601 sequence. The scores for each tested 147-bp sequence were heatmapped at the center position of that sequence. The heatmap scale indicates the local HBA score. Black arrows indicate high-HBA-score positions (124, 135, 143, 164, and 175). Red arrow indicates the dyad position of the *in vitro* reconstituted nucleosome (154). The region evaluated in (**B**) is indicated as a horizontal gray bar under the X-axis. (**B**) Local HBA scores at (red) and around (black) the *in vitro* translational position for each segment are plotted in columns. The scores for the M segment are magnified to the left to show the indices on the X-axis. The relative positions −4, 0, and +4 correspond to nucleotide positions 150, 154, and 158 of the 282-bp sequence. The location of the R3 element is indicated at the top. (**C**) Schematic representation of shifting of translational positions in (**B**). Nucleotide positions relative to the dyad of the central translated position (i=154) are indicated. Only three (−4, 0, and +4) of the nine positions evaluated in (**B**) are shown.

Next, we focused on the 147-bp Widom 601 sequence, of which the dyad base is located at nucleotide position 154 of the original sequence (**Figure 5A**, red arrow). We chose this sequence because its derivatives are well-characterized in reconstitution studies, as it forms a homogeneous nucleosome, and because the left and right halves exhibit different features [44–48]. In agreement with previous transcription studies [44, 45, 49], local HBA scores for the 147-bp sequence were highest for segments D and E, both of which share SHL −2.5, in which the high-affinity R3 element is located (**Figure 5B**, red dots). The region covered by segments D and E (SHL −3.5 to −1.5) is also known for its very strong histone-DNA interaction [46]. These data indicate that local HBA scores are useful for evaluating the suitability of nucleosomal subsegments for histone-DNA contacts.

When the translational position was shifted along the 282-bp sequence from nucleotide position 154 to surrounding nucleotide positions ranging from 150 to 158 (**Figure 5C**, −4 to +4 with respect to 154), the local HBA scores for segments B through I changed dramatically (**Figure 5B**, black dots). However, such shifting resulted in modest changes at segments J through M, with their scores remaining at relatively high levels. These observations suggest that segments J through M in the right-half sequence are generally suitable for nucleosome formation, which may also be true even when the translational position is shifted by several base pairs. In other words, segments J through M do not seem to have intra-nucleosomal disfavored elements that strongly limit the rotational setting. This nature might account for the observed poor crystal development or lower homogeneity [46], as this ‘slippery’ sequence could disrupt the uniformity of histone-DNA interactions.

### Local HBA scores for modified nucleosome-forming sequences

The above testing with the sequence previously used in reconstitution studies suggested that calculating local HBA scores would enhance understanding of how DNA sequences characterize the nucleosomes formed *in vitro*. This prompted us to examine whether we could apply local HBA calculations to the evaluation of intracellular nucleosome formation. González et al. reported that sequence-based determinants of nucleosome positioning are dispersed across nucleosomes [42]. One gene that they thoroughly investigated was the fission yeast gene, *ura4*^+^. This gene has six nucleosomes (designated +1 to +6) that overlap the coding sequence, five of which were predictable with the chemical map–based dHMM (**Additional file 1: Figure S3**, nuCpos). Although the remaining nucleosome (+5) was not clearly observed in the predicted occupancy plot, there was one position with a relatively high HBA score that reasonably corresponded to the *in vivo* nucleosome (marked with an asterisk in the figure). A heatmap of local HBA scores for this gene suggested that there is an element disfavored for nucleosome formation near the +5 nucleosome (**Figure 6**, WT, see the blueish slanted line spanning segments A through M around dyad position 600-750). Thus, this element may have caused the lower predicted occupancy of the +5 nucleosome.

**Figure 6.**
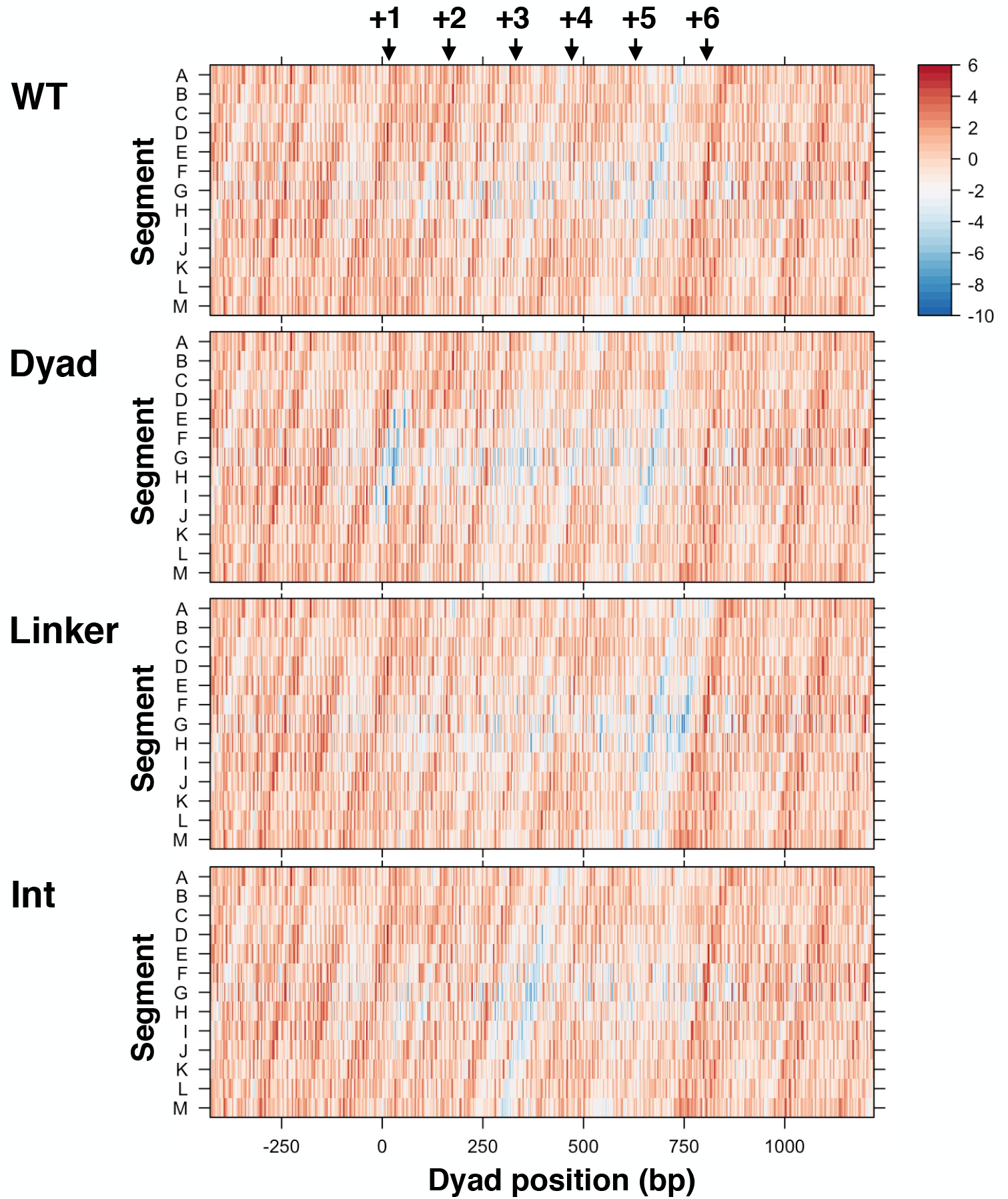
Local HBA analysis of modified nucleosome-forming sequences. Chemical map–based local HBA scores along the original and modified fission yeast *ura4*^+^gene. The scores for each tested 147-bp sequence were heatmapped at the center position of that sequence. The heatmap scale indicates the local HBA score. Black arrows indicate the nucleosome centers (nucleotide positions +17, +164, +332, +470, +629, and +806) determined by MNase-seq [42].

González et al. generated three modified sequences, namely Dyad, Linker, and Int, for the *ura4*^+^ gene and studied nucleosome positioning on these sequences *in vivo* [42]. In Dyad and Linker, 51-bp sequences centered to each nucleosome dyad and each linker, respectively, were replaced with artificial randomized sequences of the same length. In the sequence Int, two intra-nucleosomal sequences of each nucleosome (positions from −51 to −24 and 24 to 51) were replaced. According to their report, the occupancy of the +1 nucleosome was markedly decreased in the Dyad sequence, that of the +2 nucleosome was slightly increased in the Dyad and Linker sequences, and the position of the +3 nucleosome was shifted in the Int sequence [42]. However, why these changes selectively occur on specific nucleosomes *in vivo* remains unclear due to a lack of appropriate analytical methods.

A heatmap of local HBA scores for the Dyad sequence clearly showed that the replacements in this sequence caused a marked decrease in local HBA scores only around the position of the +1 nucleosome (**Figure 6**, Dyad, blue blots not present in the WT map). In contrast, the 51-bp replacements at other nucleosomal centers did not appear to dramatically change their suitability for nucleosome formation. As to the +2 nucleosome, we did not observe any supportive signatures that could explain why the Dyad and Linker replacements slightly increased its occupancy (**Figure 6**, Dyad and Linker). We observed that the Int replacements primarily affected local HBA scores around the position of the +3 nucleosome (**Figure 6**, Int, see the blueish slanted line spanning segments A through M). This suggests that the mutation specifically affects the +3 nucleosome positioning and triggers a shift of it to a more suitable position *in vivo*. Giving these results, local HBA score calculation for nucleosomal subsegments is valuable for *in silico* assessment of modified nucleosome-forming sequences.

### Chemical map–based prediction of nucleosome positioning in mice

The observation that the chemical-map–based models successfully improved the prediction accuracy of nucleosomes in both yeasts suggested that these approaches would be useful for mice. We therefore examined nucleosome positioning in the mouse genome using the MNase-seq–based and chemical map–based models. Matching of the predicted nucleosomes on the Viterbi paths at a 2-bp resolution demonstrated the better performance of the chemical map– based model (**Figure 7A**). In the MNase-seq–based prediction, 1.4% and 11.0% of predicted nucleosomes matched unique and redundant nucleosomes, respectively. In contrast, the chemical map–based prediction scored 10.0% and 41.1%, respectively. Thus, the chemical map–based model is more suitable for prediction of nucleosome positioning with respect to the genomic coordinates than the MNase-seq–based model.

**Figure 7.**
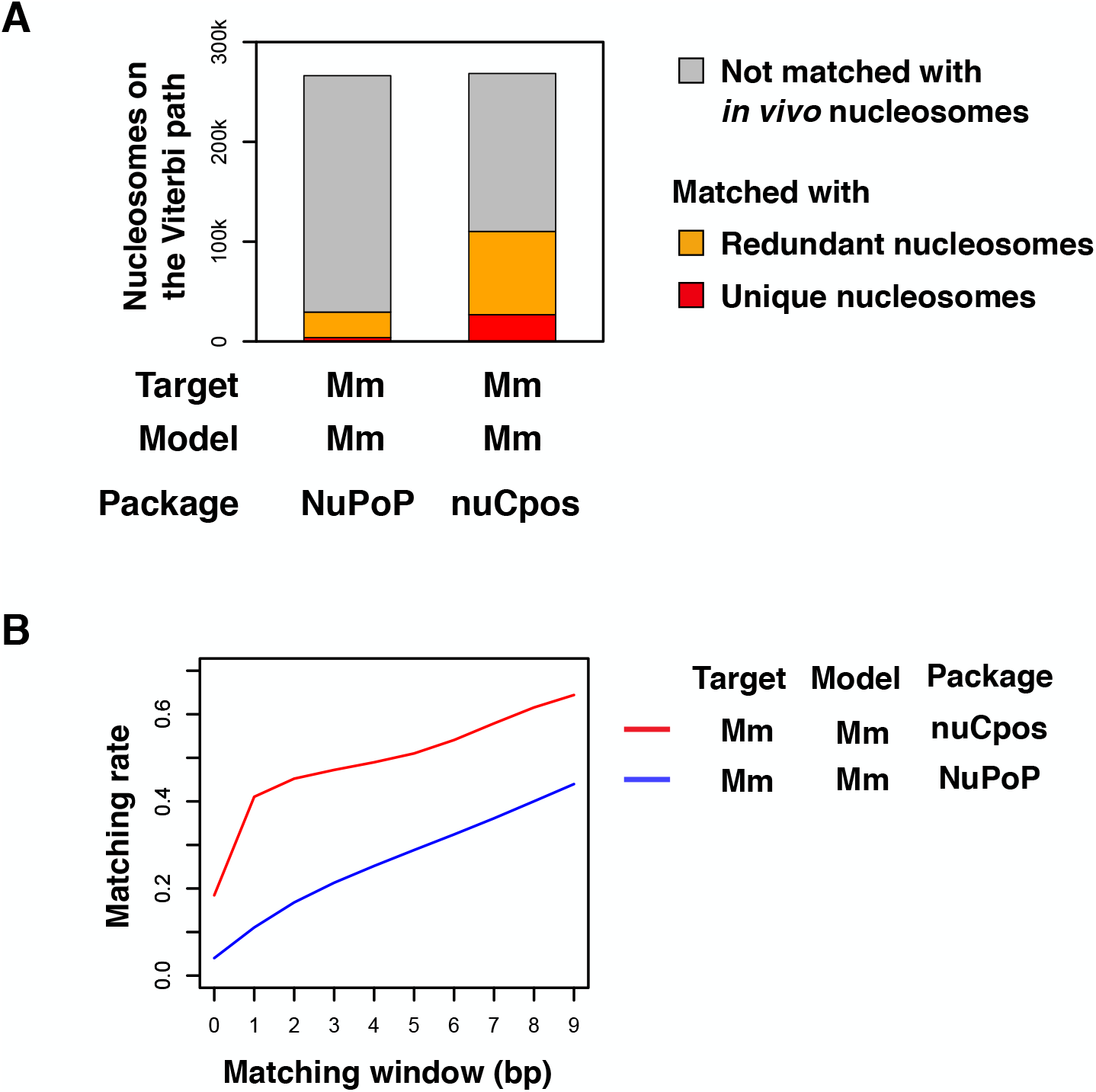
Comparison of prediction accuracy between chemical map–based and MNase-seq–based models for mice. (**A**) Matching of nucleosomes on the Viterbi paths with *in vivo* unique and redundant nucleosomes. (**B**) Matching rate between predicted and *in vivo* nucleosomes with variable matching windows. The sequence of mouse Chr19 was used for prediction.

In order to determine whether the mouse chemical map–based model could better predict the rotational setting of nucleosomes, as shown in the case of yeasts, we calculated the matching rate between predicted and *in vivo* nucleosomes with stepwise widening of the matching window (**Figure 7B**). As expected, the matching rate for MNase-seq–based prediction increased gradually as the matching window widened. In chemical map–based predictions, a dramatic increase in matching rate was observed when the matching window was widened from 0 to 1 bp. However, the matching rate also increased gradually as the matching window was widened further (See **Figures 1B and 7B** for comparison between species). The shapes of the matching rate curves suggested the chemical map–based model still distinguished rotational settings to some extent. However, the mouse chemical model did not appear to be as good at discriminating rotationally mispredicted nucleosomes compared with the yeast models.

## Discussion

We developed a chemical map–based computational method to predict nucleosome positioning and assessed the prediction results. Training dHMMs with chemical maps improved the prediction accuracy of nucleosome locations with respect to the genomic coordinates in budding yeast, fission yeast, and the house mouse, *Mus musculus* (**Figures 1 and 7**). A total of 41-54% of the predicted nucleosomes on the Viterbi paths matched those of *in vivo* nucleosomes at 2-bp resolution. Genome- and locus-level evaluations showed that the software successfully predicted the positions of *in vivo* nucleosomes (**Figures 3 and 4, and Additional file 1: Figures S2 and S3**). The *in vivo* nucleosomes generally had high HBA scores at reasonably near genomic positions. Furthermore, we demonstrated that perturbation of nucleosome positioning associated with genetic alterations could also be predicted (**Figures 4 and 6, and Additional file 1: Figure S4**). Thus, we propose that the software nuCpos can be used for predicting nucleosome positions and also for engineering of nucleosome-forming sequences.

As expected, the use of chemical maps in prediction led to strict recognition of rotational settings, which was demonstrated with *in vitro* and *in vivo* nucleosome-forming sequences (**Figures 1, 2 and 7**). Indeed, chemical map–based HBA scores for the original Widom 601 sequence indicated that the 147-bp sequence centered at nucleotide position 154 was suitable for nucleosome formation (**Figure 2**). The *in vitro* nucleosome positions formed on natural sequences were also predictable at a rotational level (**Additional file 1: Figure S1**); however, the HBA amplitude along these sequences was smaller than along the *in vitro*–optimized Widom sequence. In contrast to the chemical map–based models, the MNase-seq–based models largely failed to recognize rotational settings (**Figures 1, 2 and 7**). This may have been due to the A/T preference of MNase and the difficulty of dyad base calling in the construction of MNase-seq–based nucleosome maps [7, 12]. Indeed, the MNase-map–based model simply predicted A/T-rich regions as nucleosome-depleted regions (**Figure 3 and Additional file 1: Figure S3**). Therefore, instead of lower-resolution MNase-seq–based maps, base-pair-resolution chemical maps should be used in the development of bioinformatics methods to obtain better prediction results.

In the field of synthetic biology, the MNase-seq–based method NuPoP has contributed to the design of synthetic promoters and terminators [29, 30]. However, some studies reported that the histone binding affinity and predicted nucleosome occupancy output by NuPoP do not correlate with synthetic promoter and terminator activity [34, 35]. Differences in the DNA sequences of the tested synthetic elements may of course account for this discrepancy, as discussed elsewhere [34, 35]. Clearly, the A/T frequency does affect the prediction results of NuPoP (**Figure 3 and Additional file 1: Figure S3**). As demonstrated in the Results section, the chemical map–based method is more accurate and not affected by the A/T frequency. Thus, we expect that nuCpos is applicable to prediction-based engineering and more effective for synthesizing functional elements. Similarly, nuCpos can also be used to examine whether DNA sequences of interest are suitable for nucleosome formation, as previously done with NuPoP [37, 38].

Dividing 147-bp nucleosomal DNA into subsegments and calculating subsegment local HBA scores revealed unexpected landscapes of the tested nucleosomes. Our study demonstrated that the Widom sequence had segments that are both favored and disfavored for nucleosome formation (**Figure 5A**). Importantly, segments with high local HBA scores in the Widom 601 sequence matched the high-affinity histone-contacting regions (**Figure 5B**). Furthermore, the Widom sequence had segments exhibiting the highest local HBA scores at the *in vitro* rotational settings (**Figure 5B**). In contrast to the left-half sequence, the right-half sequence contained segments exhibiting relatively high local HBA scores that did not change markedly with different rotational settings. Thus, these sequence-specific features may account for the different behavior of each half in *in vitro* experiments [44–48]. Collectively, our data indicate that local HBA scores for given sequences provide insights that enhance understanding of nucleosome features.

Locus-level assessments of prediction results suggested that software users should not place undue confidence in the predicted nucleosome occupancy output from chemical map–based models. Indeed, there was a misplaced nucleosome that occupied the replication origin *ARS1* (**Figure 3**). We speculate that these discrepancies were due primarily to the fact that the models do not consider involvement of functional DNA elements or transacting factors that regulate nucleosome formation at specific genomic locations. In other words, chromatin regions highlighted by the differences in nucleosome occupancy between prediction and *in vivo* observations could be the targets of future investigations aiming to uncover region-specific regulatory mechanisms.

We demonstrated that the prediction accuracy of the chemical map– based mouse model (41%) was apparently lower than that of yeast models (over 50%) (**Figures 1 and 7**). A disadvantage of mouse prediction is that the chemical map probably contains substantial numbers of pseudo-positive nucleosomes, which could not be omitted due to the lower cleavage depth than yeasts [22]. As a consequence, the density of nucleosomes on the mouse genome is one nucleosome per 3 bp, which is about 10 times higher than yeast values (see **Methods**). Thus, we speculate that the accuracy of mouse nucleosome positioning prediction can be increased if other chemical maps of higher quality are produced and used for model construction. In human genomics, the relationship between various phenotypes and genetic variants has been explored [50–53]. However, the effects of genetic variants on nucleosome formation in this regard remain to be studied. Now that our study showed that insertions and replacements that disrupt nucleosomes in yeast cells can be predictable *in silico*, it is possible that nucleosome-disrupting human mutations can be found with the aid of chemical map–based predictions.

## Conclusions

In this study, we demonstrated that the accuracy of dHMM-based nucleosome positioning prediction can be substantially increased by using base-pair-resolution nucleosome maps for model construction. Our prediction results suggest that chemical map–based models are useful for predicting nucleosome positioning in wild-type and modified sequences at the locus level. We also demonstrated that strong histone-DNA contacts in a nucleosome and their rotational settings can be predicted. Furthermore, as another advantage of bioinformatics methods, our models indicate that the commonly used Widom sequence contains subnucleosomal segments that are disfavored, or statistically very rare in *in vivo* nucleosomes, at their nucleosomal positions in shifted rotational settings. We expect that our prediction method will provide further insights that will enhance understanding of nucleosome-based epigenetic regulation.

## Methods

### Software and data sets

Most analyses were performed in the GNU R environment (https://www.r-project.org, ver. 3.6.1). R packages were obtained from CRAN (https://cran.r-project.org/) and Bioconductor (http://bioconductor.org/). Chemical maps for budding yeast, fission yeast, and house mouse (*Mus musculus*) embryonic stem cells [17, 19, 22] were used for model construction and testing the prediction accuracy. The budding yeast’s chemical map in the sacCer2 coordinate was lifted over to the sacCer3 coordinate, as described elsewhere [27]. The number of unique and redundant nucleosomes in the budding yeast genome was 67,548 and 344,709, respectively; fission yeast, 75,828 and 425,653; mice, 10,677,016 and 850,701,275. Reference genomes of budding yeast (R64-1-1), fission yeast (ASM294v2), and mice (mm9/NCBIM37.67) were used. The original 282-bp Widom 601 sequence [10], the 485-bp MMTV 3’–LTR sequence [54] and the somatic 5S RNA gene of *Xenopus borealis* [55] were used as test sequences.

### Construction of the nuCpos package

The codes for the construction of parameters used in the nuCpos package (ver. 1.2.0) are available online (**https://doi.org/10.5281/zenodo.3362065**). Genomic regions covered with the 147-bp non-redundant (unique) chemically mapped nucleosomes and uncovered were defined as nucleosome and linker regions, respectively. DNA sequences of these regions were used to construct parameters that were transferred to internal Fortran programs. Nucleosomes of which dyads were located within 73 bp of the chromosomal ends were omitted. For construction of the mouse model, hard-masked genomic sequences were used, and nucleosome and linker regions containing N were omitted before parameter construction to avoid potential prediction bias caused by repeat elements. In total, 67,538 nucleosome regions and 50,622 linker regions were obtained for the budding yeast genome (sacCer3); fission yeast (ASM294v2), 75,826 nucleosome and 46,557 linker regions; mice (mm9), 4,147,972 nucleosome and 2,484,347 linker regions.

We developed an R function designated *predNuCpos*, which predicts nucleosome positioning based on a dHMM, as previously proposed by Xi et al. [16]. Like its ancestral function *predNuPoP* in the NuPoP package (https://doi.org/doi:10.18129/B9.bioc.NuPoP, ver. 1.34.0), *predNuCpos* receives a DNA sequence of any length, invokes an internal Fortran program, and outputs the prediction result either in the working directory or in the working environment of R. In *predNuCpos*, construction of the dHMM is based on chemical maps, as described below.

Parameters used in the *predNuCpos* function were constructed according to the NuPoP paper [16] using the functionalities of the Biostrings package (https://doi.org/doi:10.18129/B9.bioc.Biostrings, ver. 2.52.0). The parameters were as follows: *freqL*, one-base frequencies for linker regions; *tranL*, *tranL2*, *tranL3*, and *tranL4*, First- to 4th-order transition probabilities for linker regions, respectively; *freqN4*, four-base frequencies at the first four nucleotide positions of nucleosome regions; *tranN4*, time-dependent 4th-order transition probabilities for nucleosome regions; *Pd*, linker length distribution that ranges from 1 to 500 bp. Linker sequences of 7-500 bp in length were used for linker model construction, as described elsewhere [16]. *freqL* and *freqN4* were obtained using the *oligonucleotideFrequency* function of Biostrings; *tranL*, *tranL2*, *tranL3*, *tranL4*, and *tranN4* were obtained using the *oligonucleotideTransitions* function of Biostrings. Moving average smoothing using the *SMA* function of the TTR package (https://cran.r-project.org/package=TTR, ver. 0.23-4) at a 3-bp window was applied to the 4th-order transition probability parameter *tranN4* and to the linker length distribution parameter *Pd*. The parameters used in *predNuCpos* were also used in another function, *mutNuCpos*, which predicts the effect of genetic alterations on nucleosome positioning.

Xi et al. proposed the HBA score [16], which is also referred to as the ‘nucleosome affinity score’, as the log likelihood ratio of the probability for a given 147-bp sequence to be a nucleosome versus a linker. According to their definition, the HBA score for the 147-bp region *x* centering at position *i* (*ai*) on a given genomic sequence is,

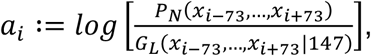

where *P*_*N*_ and *G*_*L*_ represent the probability of observing the 147-bp sequence as a nucleosome or a linker, respectively [16]. The probability of being a nucleosome is calculated by referring to the parameters *freqN4* and *tranN4*, which are derived from nucleosomal DNA sequences. Similarly, calculation of the probability of being a linker is based on linker DNA sequences. As nucleosomal and linker sequences do not overlap in terms of their genomic coordinates, negativity of HBA does not directly mean that the tested sequence is inappropriate for nucleosome formation. The *predNuCpos* function calculates chemical map–based HBA scores along the input sequence and outputs them as raw values as its default behavior. We developed an independent function designated *HBA*, which only calculates the HBA score for a given 147-bp sequence. The *HBA* function uses the abovementioned chemical parameters for *predNuCpos*: *freqL*, *tranL*, *tranL2*, *tranL3*, *tranL4*, *freqN4*, and *tranN4*.

We defined 13 overlapping nucleosomal subsegments, A through M, and developed a function designated *localHBA* that calculates “local” HBA scores for each segment. Segment A corresponds to nucleosomal nucleotide positions 1-21; B, 12-31; C, 22-42; D, 33-52; E, 43-63; F, 54-73; G, 64-84; H, 75-94; I, 85-105; J, 96-115; K, 106-126; L, 117-136; and M, 127-147. Similar to the calculation of HBA [16], the local HBA score for segment A of the 147-bp potential nucleosomal region *x* centering at position *i* (*l_i_*) on a given genomic sequence is calculated as,

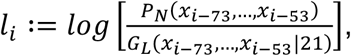

where the probabilities of observing the 21-bp sequence as segment A of a nucleosome and a linker are calculated. Local HBA scores for the other segments are calculated in the same way, except that the considered nucleotide positions are set appropriately. At the implementation level, four-base frequency values for the first four nucleotide positions of each segment were prepared: *freqN4SA* corresponds to nucleosome positions 1-4; *freqN4SB*, 12-15; *freqN4SC*, 22-25; *freqN4SD*, 33-36; *freqN4SE*, 43-46; *freqN4SF*, 54-57; *freqN4SG*, 64-67; *freqN4SH*, 75-78, *freqN4SI*, 85-88; *freqN4SJ*, 96-99; *freqN4SK*, 106-109; *freqN4SL*, 117-120; and *freqN4SM*, 127-130. These four-base frequency values were used to calculate the probability of the segment as that part of nucleosomal DNA as done for HBA calculations [16].

### Prediction of nucleosome positioning with nuCpos and NuPoP and evaluation of the prediction results

The codes for prediction, evaluation, and figure drawing used in this study are available online (https://doi.org/10.5281/zenodo.4083950). Unmasked reference genomes for the budding yeast, fission yeast, and house mouse (*Mus musculus*) were used as prediction target sequences. For the mouse experiments, only the 19th chromosome (Chr19) was used for prediction because its length (61,342,430 bp) was sufficient for evaluation, at approximately five-times longer than the total length of yeast chromosomes (budding yeast, 12,071,326 bp; fission yeast, 12,571,820 bp). Mouse Chr19 contained 249,210 unique and 19,965,481 redundant nucleosomes that had been determined based on histone H4 S47C-dependent cleavage [22]. The density of redundant nucleosomes on mouse Chr19 was one per 3.07 bp. With this density, one in three randomly located nucleosomes in the chromosome could be counted as a truly predicted nucleosome. Thus, we selected 2,044,747 nucleosomes with the highest NCP scores as true redundant nucleosomes for the matching experiment. This selection yielded a density of one nucleosome per 30.0 bp, which was comparable to that of budding yeast and fission yeast (one nucleosome per 35.0 bp and 29.5 bp, respectively).

The *predNuPoP* function of NuPoP and the *predNuCpos* function of nuCpos were used for dHMM-based predictions. In order to predict using budding yeast models, the species arguments were set as “7” for *predNuPoP* and “sc” for *predNuCpos*. Similarly, fission yeast (“9” and “sp”) and mouse (“2” and “mm”) models were specified. We defined any nucleosomes on the Viterbi path as predicted nucleosomes. Matching of predicted nucleosomes using *in vivo* chemical nucleosomes was done by widening the genomic coordinates of *in vivo* nucleosomes to both sides in a step-wise manner from zero to nine base pairs. For instance, at 1-bp resolution, a predicted nucleosome was regarded as “matched” when the genomic coordinate of its dyad was equal to that of the nearest *in vivo* nucleosome. If the dyad of the predicted nucleosome was 1 bp away from that of the nearest *in vivo* nucleosome, it was regarded as “not matched.”

Chemical map–based HBA scores were calculated using the *predNuCpos* function or the *HBA* function of nuCpos, which yield the same scores. The species argument was set as “sc” for budding yeast sequences and sequences of *in vitro* reconstitution studies; for the fission yeast sequences, “sp.” The *predNuPoP* function of NuPoP smooths raw HBA scores using a moving average of a 55-bp window before outputting the processed HBA scores. This prevents software users from examining the actual HBA score for each genomic coordinate to determine if the 147-bp sequence of interest is suitable for nucleosome formation. To overcome this problem, non-smoothed MNase-seq–based HBA scores were calculated using in-house functions that utilize the parameters for the *predNuPoP* function of NuPoP. For comparison of *in vivo* nucleosomes, 5,542 sets of −1 and +1 nucleosomes determined by Chereji et al. [12] were used for HBA calculations. The 347-bp sequences around these nucleosomes (from −173 to +173 nucleotide positions with respect to the dyads) were extracted from the budding yeast genome. The HBA score was calculated for each possible 147-bp segment along the sequences. A/T-frequency was calculated using the *letterFrequency* function of Biostrings for each 147-bp sequence. HBA and A/T-frequency scores were averaged at each coordinate with respect to the −1 or +1 nucleosomes. Chemical map–based local HBA scores were calculated using the *localHBA* function of nuCpos. Heatmaps of local HBA scores were drawn using the *levelplot* function of the rasterVis package (https://cran.r-project.org/package=rasterVis, ver. 0.46).

## List of abbreviations

dHMM: duration hidden Markov model
HBA: histone binding affinity
LTR: long terminal repeat
MMTV: mouse mammary tumor virus
MNase: micrococcal nuclease
NCP: nucleosome core particle
NDR: nucleosome depleted region
SELEX: Systematic Evolution of Ligands by EXponential enrichment
SHL: superhelical location

## Declarations

### Ethics approval and consent to participate

Not applicable.

### Consent for publication

Not applicable.

### Availability of data and materials

The datasets generated and analyzed during the current study are available online (see **Methods** for the URLs).

### Authors’ contributions

H.K. designed the study, created the software, performed the experiments, and was a major contributor in writing the manuscript. T.U. and M.S. were involved in study design, data validation, and manuscript writing. All authors read and approved the final manuscript.

## Acknowledgments

The authors thank Dr. K. Ura for providing the dinucleosome-forming sequence and Dr. Y. Ichikawa for confirming the DNA sequences. The authors also thank Drs. H. Kimura, H. Kurumizaka, J. Nakayama, and Y. Ohkawa for their helpful comments and encouragement.

## Supplementary Figure Legends

**Additional file 1: Figure S1. HBA scores along *in vitro* nucleosome-forming sequences.** HBA scores along nucleosome-forming sequences were calculated using chemical map–based and MNase-seq–based budding yeast models. Sequences analyzed were the 3’-LTR of MMTV (**A**) and the *Xenopus borealis* 5S rDNA dinucleosome-forming sequence (**B**). The scores were normalized by subtracting the mean value from the raw value on a per sequence basis. The HBA score for a given 147-bp nucleosome sequence was assigned to its dyad position. Orange vertical lines indicate the dyad positions of *in vitro* reconstituted nucleosomes: nucleotide positions 139 and 335 (**A**) and 106 and 303 (**B**). Asterisks indicate high-scoring HBA positions around the *in vitro* positioning sites. Note that for the 5S rDNA sequence, two identical sequences are joined at position 230, causing a difference in HBA scores around the left sides of the two *in vitro* nucleosome positions.

**Additional file 1: Figure S2. Magnified view of prediction results for *TRP1ARS1***. Prediction results for the budding yeast *TRP1ARS1* mini-chromosome output by nuCpos and NuPoP (**Figure 3C**) were magnified, being centered at nucleosome II.

**Additional file 1: Figure S3. Prediction results for the fission yeast *ura4*^+^ gene.** Schematic representation of *in vivo* nucleosome positioning is shown above the plots. Nucleosomes numbered +1 to +6 are indicated as ovals. The top two panels show the prediction results output by nuCpos, whereas the next two panels show NuPoP results. The upper panel in each set shows predicted occupancy of nucleosomes (Occup., gray polygons) and probability that the tested 147-bp sequences are in the nucleosome state (P-dyad, blue vertical lines). Lower panels show HBA values for the tested 147-bp sequences calculated using the indicated models. The very bottom panel shows the A/T-frequency for the tested 147-bp sequences. Horizontal lines at the bottom of each plot indicate the 5’- and 3’-untranslated regions (colored in gray, nucleotide positions −151 to −1 and +795 to +986, respectively) and the protein-coding region (red, 0 to +794) of the *ura4*^+^ gene. Inverted triangles indicate the nucleosome centers determined by MNase-seq [42].

**Additional file 1: Figure S4. Prediction of the effects of repeat insertion in the fourth nucleosome of the *TALS* minichromosome.** Chemical map–based HBA scores and predicted nucleosomal occupancy were calculated for original and modified *TALS* sequences [43]. The sequences were centered at the dyad position of the fourth nucleosome (0 bp), in which the indicated repeat sequences were inserted. Note: the position for the fourth nucleosome was predicted to shift to the right. Horizontal lines at the bottom of each plot indicate the *α2*-operator (purple) and the insert (red). hTEL stands for human telomeric repeat (5’-TTAGGG-3’); SI-A, sequence isomer-A (5’-TGTAGG-3’); SI-B, sequence isomer-B (5’-TGTGAG-3’). When telomeric DNA fragments are inserted at the center of the fourth nucleosome, shorter fragments (hTEL2 and hTEL4) were predicted to not affect nucleosome positioning. In contrast, longer fragments (hTEL12 and hTEL29) were predicted to severely compromise nucleosome formation. In addition, sequence isomers of the telomeric repeat (SI-A6, SI-A12, SI-B6, and SI-B12) were predicted to not inhibit nucleosome formation. These prediction results agreed with previous *in vivo* observations [43]. The hTEL6 insertion, which causes nucleosome depletion *in vivo*, appeared not to affect nucleosome formation in the prediction, suggesting a limitation of the prediction.

**Additional file 1: Figure S5. Thirteen overlapping nucleosomal DNA subsegments for local HBA calculation.** Segments A through M are colored in orange on the nucleosome structure 1AOI [1]. The structures were drawn using MacPyMOL (v1.8.4.1). The base pair length (red) and corresponding superhelical locations (black) of each segment are indicated. The color of the right-half segments (H-M) is faded because they are behind the front helix.

