## Supplemental Figures for "Chemical map–based prediction of nucleosome positioning using the Bioconductor package nuCpos"

Hiroaki Kato[1], Mitsuhiro Shimizu[2] and Takeshi Urano[1]

[1]Department of Biochemistry, Shimane University School of Medicine, Izumo, Shimane 693-8501, Japan

[2]Department of Chemistry, Graduate School of Science and Engineering, Program in Chemistry and Life Science, School of Science and Engineering, Meisei University, Hino, Tokyo 191-8506, Japan

### Additional file 1: Figure S1

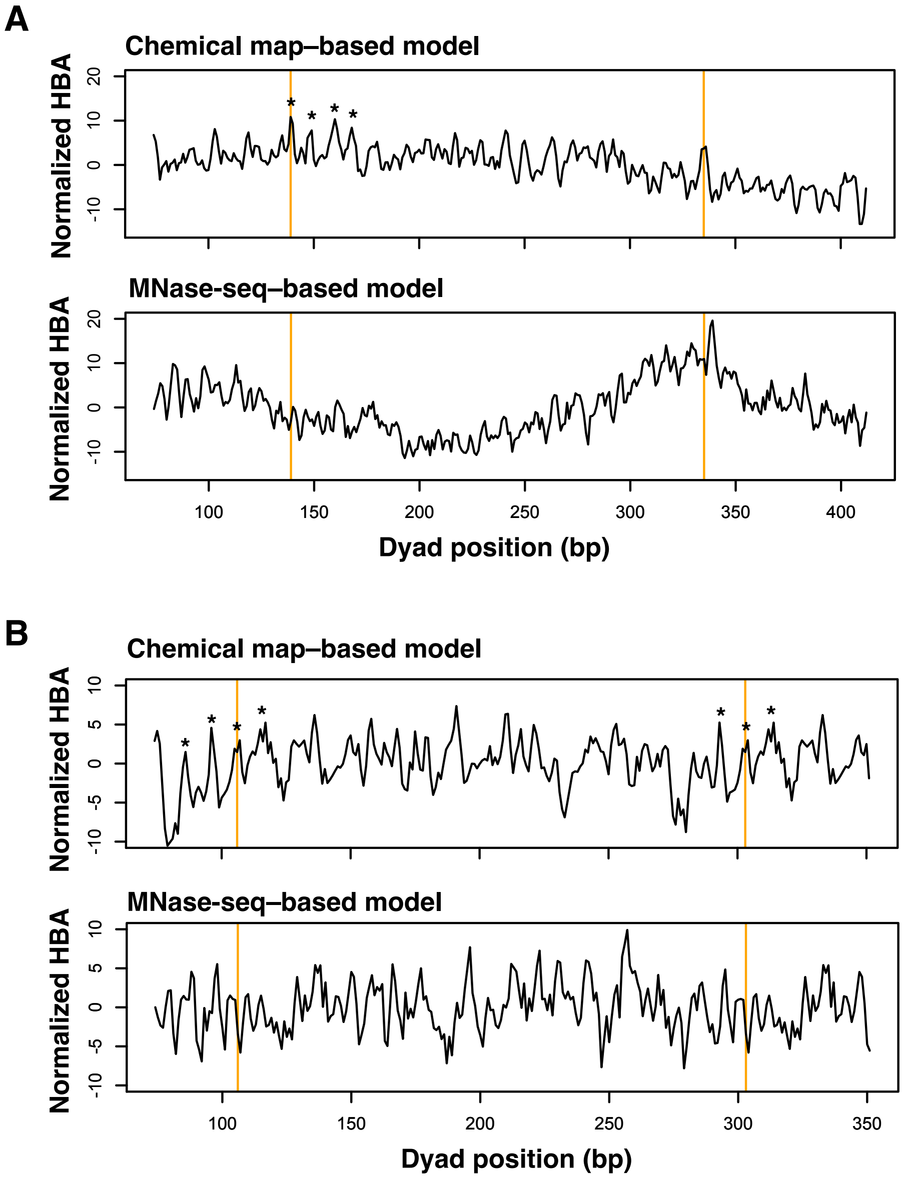

### Additional file 1: Figure S2

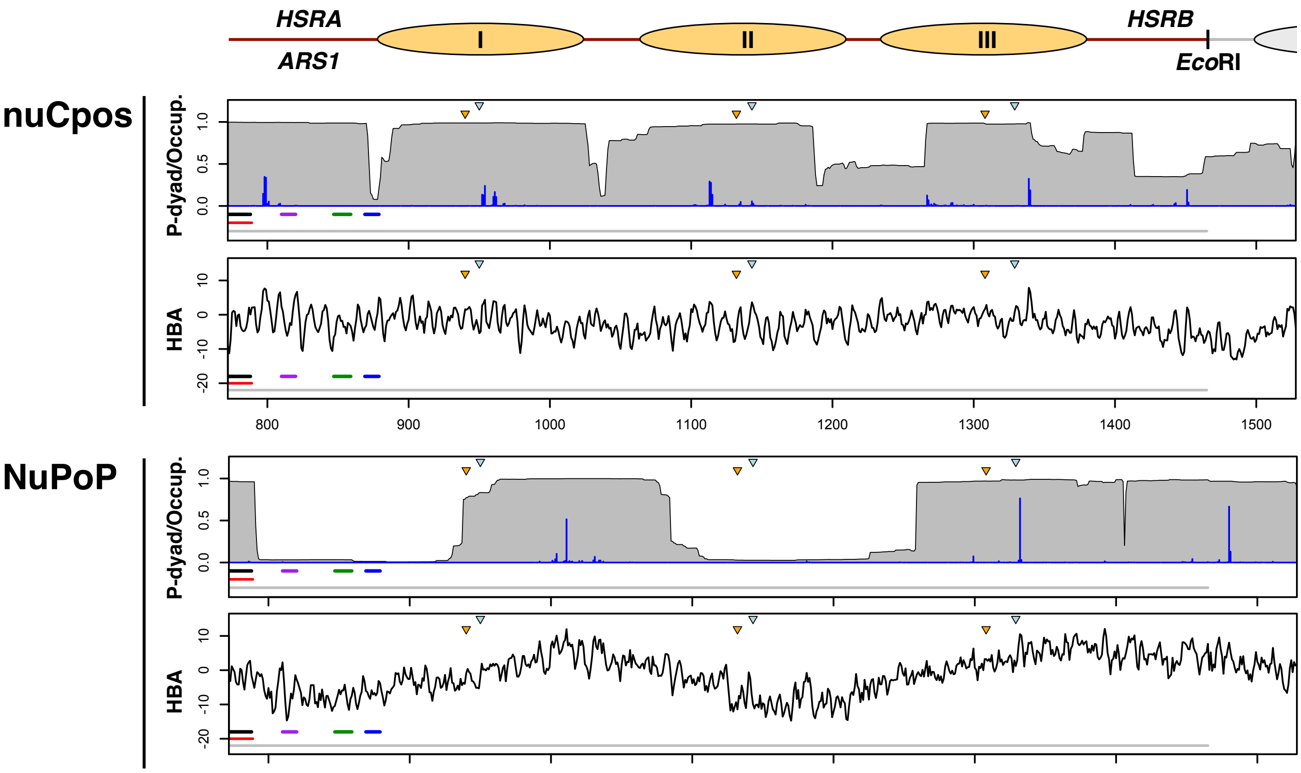

#### Additional file 1: Figure S3

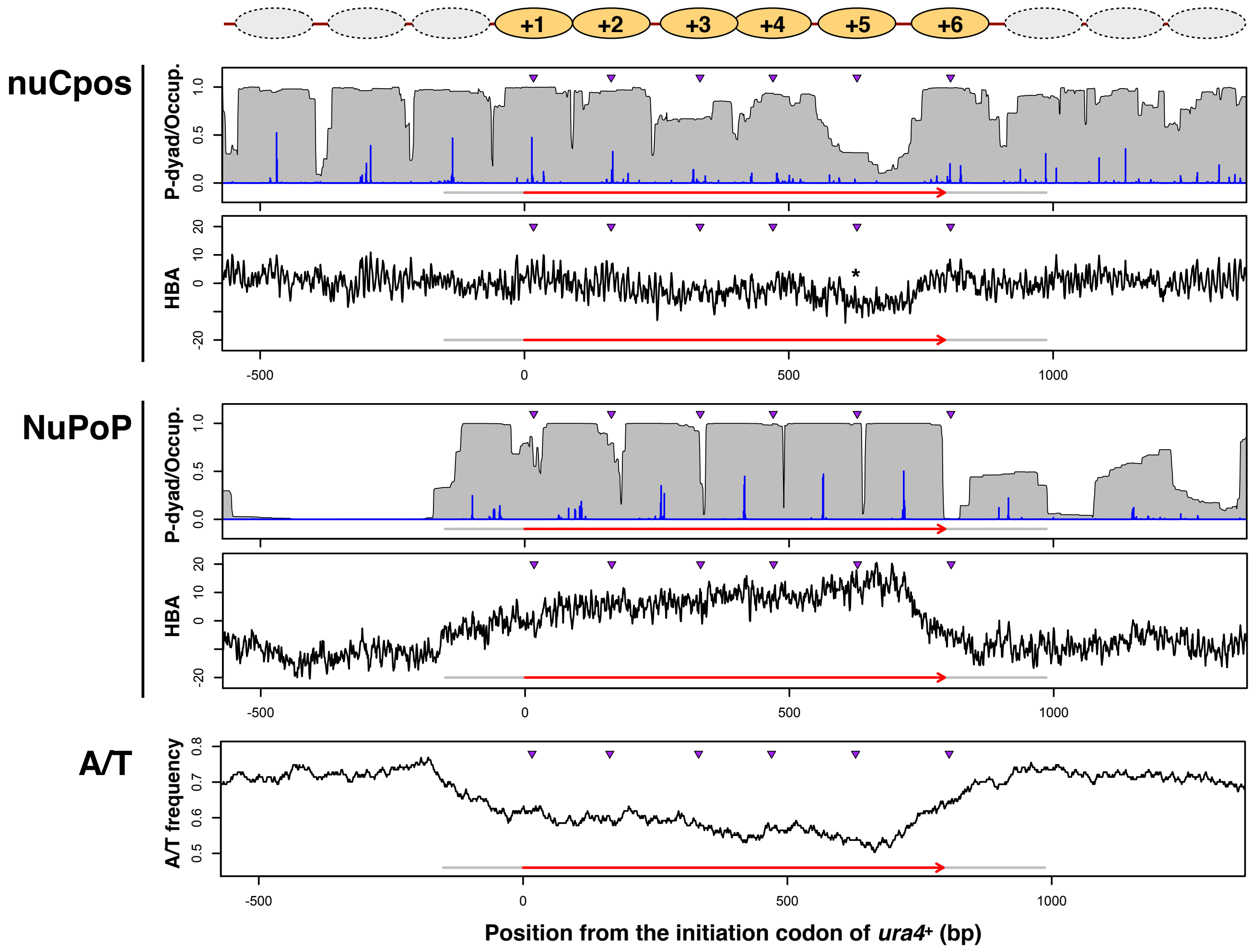

#### Additional file 1: Figure S4

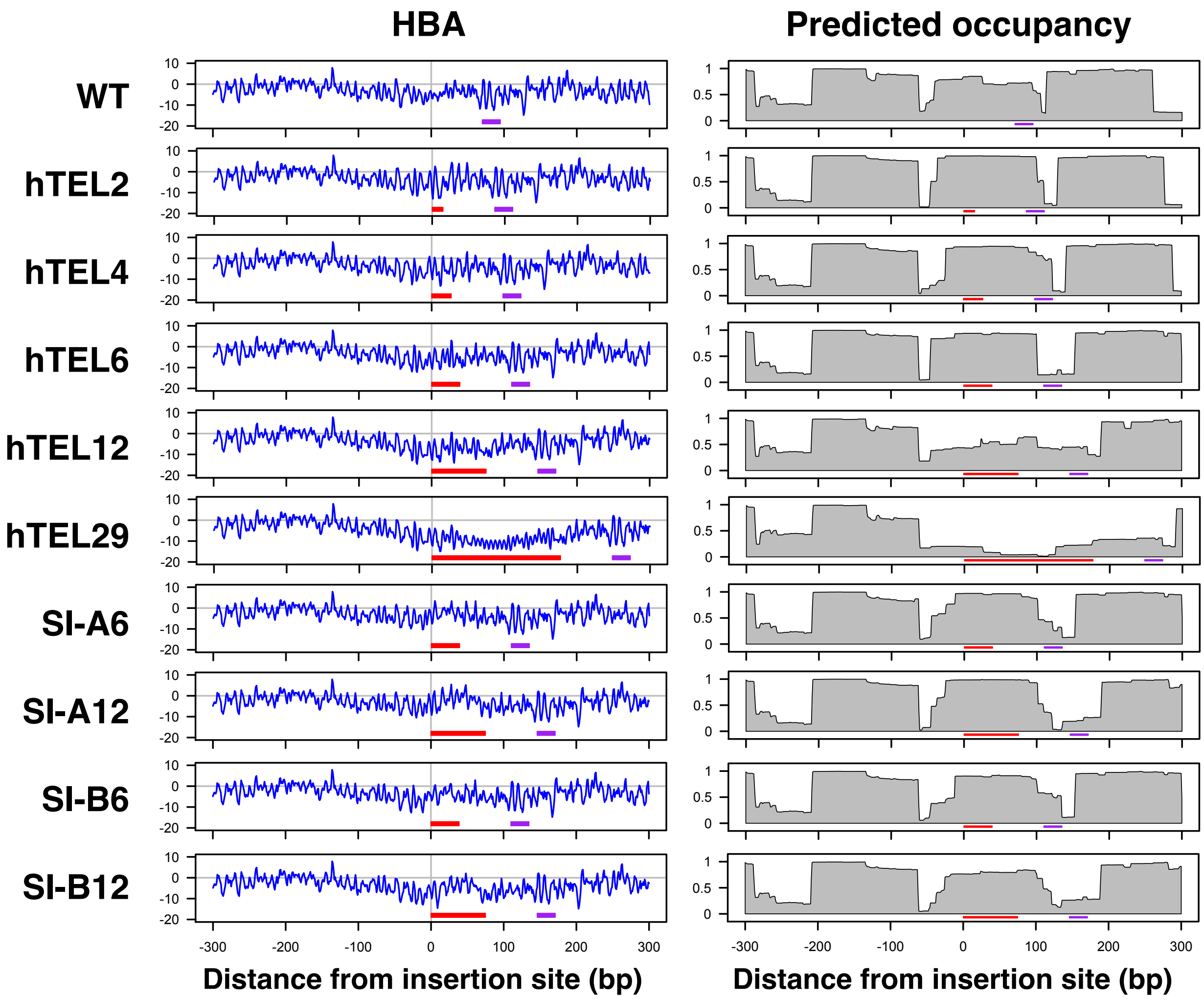

#### Additional file 1: Figure S5

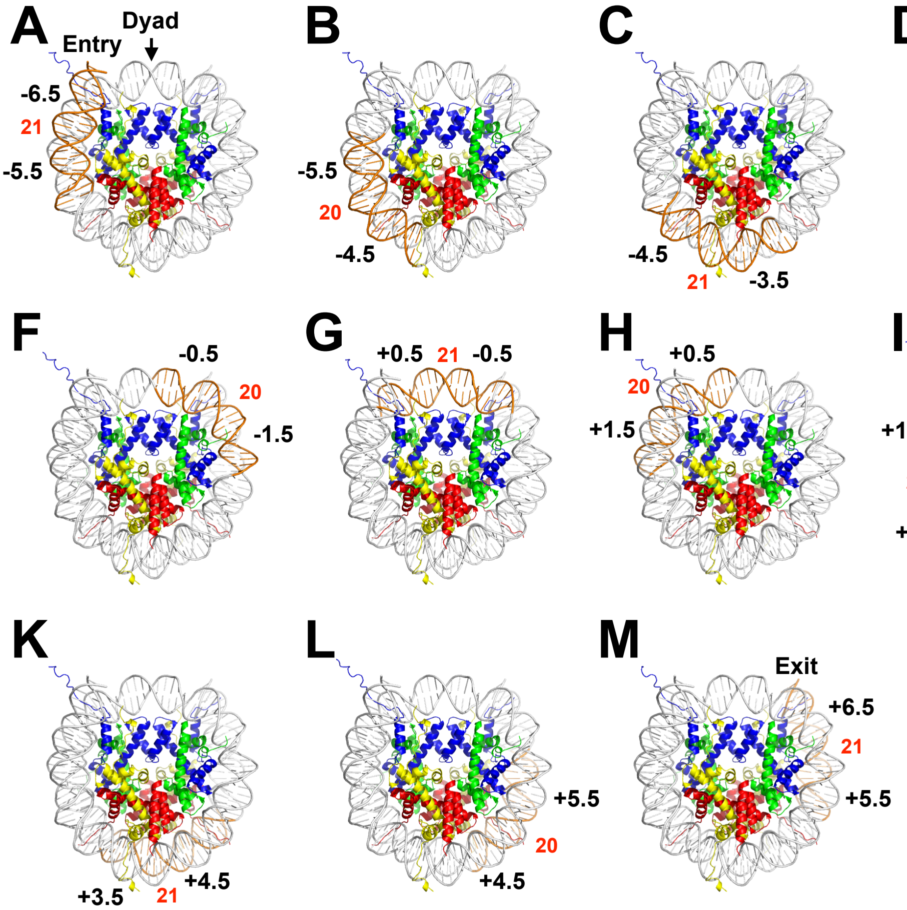
